# Contrast-Free Microvascular and Functional Brain Imaging by Sparse Deconvolution of Ultrafast Power Doppler

**DOI:** 10.64898/2026.09.10.750681

**Authors:** Ge Zhang, Nicolas Zucker, Thomas Deffieux, Mathieu Pernot, Nathalie Ialy-Radio, Sophie Pezet, Mickael Tanter

## Abstract

Ultrafast power Doppler imaging combined with singular value decomposition (SVD) clutter filtering has become a standard approach for label-free microvascular ultrasound, enabling the visualization of small vessels without microbubble contrast agents. In the absence of contrast, however, SVD-filtered Doppler images remain limited by the blur of the imaging-system point spread function (PSF) and by a residual noise floor that reduces sensitivity at depth, which together hinder the resolution of fine microvasculature. Here we establish a sparse deconvolution framework to SVD-filtered ultrafast power Doppler images. Each Doppler frame is processed in two cascaded stages: a Split-Bregman optimization that solves a regularized least-squares problem combining an ℓ1 sparsity prior and a Hessian continuity prior, followed by an accelerated Richardson–Lucy deconvolution with an estimated system PSF. We first validated the framework on a simulation phantom with known ground truth, and then evaluated the framework on in vivo rat-brain plane-wave acquisitions obtained with a Verasonics Vantage system and a 15-MHz linear array. Compared to conventional SVD power Doppler, sparse deconvolution improved the resolution by around 4 and 8 times to λ/2 and λ/4, in the lateral and axial directions respectively. We further show that decomposing the Doppler signal into velocity bands before deconvolution disentangles slow and fast flow and yields a velocity-resolved microvascular map. Finally, applying the same framework to task-evoked functional ultrasound, we show that sparse deconvolution preserves the stimulus-locked cerebral-blood-volume response measured by conventional functional ultrasound while sharpening the corresponding activation map from a diffuse cortical region to discrete penetrating vessels. These results indicate that the sparsity-prior super-resolution principles established in label-free ultrafast Doppler ultrasound, and that sparse deconvolution can serve as a practical, contrast-free post-processing front-end for super-resolution microvascular and functional imaging.

## 1. Introduction

Microvascular imaging plays a central role in characterizing tissue physiology and pathology, since changes at the level of small vessels and capillaries are early indicators of stroke, tumor angiogenesis, neurodegeneration, and inflammation. Among non-invasive modalities, ultrasound stands out for its high temporal resolution, portability, and absence of ionizing radiation. Yet conventional Doppler ultrasound has long been restricted to large vessels by the limited frame rate of focused-beam imaging and by the poor sensitivity of Doppler estimators to low blood-flow velocities.

The advent of ultrafast plane-wave imaging, combined with singular value decomposition (SVD) clutter filtering, has dramatically expanded the sensitivity of ultrasound to slow blood flow and small vessels. By acquiring thousands of frames per second and exploiting the spatiotemporal coherence of tissue motion versus blood flow, SVD-based ultrafast Doppler separates blood signals from tissue clutter [1–3]. This approach has become the de facto standard for label-free microvascular ultrasound, since it enables visualization of small vessels without recourse to microbubble contrast agents and has been widely applied to functional ultrasound imaging of the brain, kidney, and tumor microenvironments [4–6].

Despite this progress, contrast-free ultrafast Doppler images suffer from two intrinsic limitations. First, the lateral and axial resolutions are bounded by the system point spread function (PSF), which for a 15-MHz linear array is on the order of 200 µm and prevents the resolution of capillary-scale microvasculature. Second, in the absence of microbubble enhancement, the blood signal is several orders of magnitude weaker than the tissue clutter, so that residual noise after SVD filtering severely degrades the visibility of deep vessels. Microbubble-based ultrasound localization microscopy (ULM) [7, 8] circumvents both limitations and reaches micrometer-scale resolution, but it requires the injection of an exogenous contrast agent, prolonged acquisitions of several minutes, and dedicated post-processing, which limits its translation to bedside or repeated longitudinal use.

In optical microscopy, similar resolution and contrast trade-offs have been addressed through computational super-resolution. Recently, Zhao et al. introduced a sparse deconvolution framework that exploits two physical priors of biological structures, namely sparsity and continuity, to push the spatial resolution of fluorescence microscopes nearly twofold without any hardware modification [9]. The method casts reconstruction as the minimization of a regularized least-squares cost combining an ℓ1 sparsity term and a Hessian continuity term, and it is followed by an iterative Richardson–Lucy deconvolution that recovers high-frequency content within the system optical transfer function (OTF). Because both sparsity and continuity are general features of fluorescence images rather than content-specific assumptions, the framework has been shown to generalize across structured-illumination, confocal, two-photon, and STED microscopes [9].

Within ultrasound itself, deconvolution and sparse signal recovery have a long history. Restoration of B-mode and Doppler images with estimated or measured PSFs has been used to sharpen tissue and vascular structure [10–12], and sparsity-promoting and compressed-sensing formulations have been applied both to beamforming and to microbubble localization in ULM, where they help resolve overlapping point sources [13, 14]. More recently, deep-learning methods have been proposed to denoise or super-resolve power-Doppler and microvascular maps [15, 16]. These methods, however, either target microbubble data, depend on learned priors and training data, or act at the channel/beamforming level. The approach pursued here is distinct in that it operates directly on the SVD-filtered power-Doppler frames produced by the standard contrast-free pipeline, requires no training data, and combines an explicit, content-agnostic pair of physical priors, which are sparsity and Hessian continuity, with a calibrated-PSF Richardson–Lucy stage.

Several contrast-free or super-resolution ultrasound strategies have likewise sought to overcome this resolution limit, yet each carries specific constraints. Ultrasound localization microscopy (ULM) attains micrometer-scale resolution but relies on spatially isolated microbubble signals and long temporal accumulation [7]. Super-resolution ultrasound using erythrocytes (SURE) exploits global peak detection of red-blood-cell scattering and therefore demands a high vessel-to-noise ratio [17]. Null subtraction imaging (NSI) refines the effective point spread function but requires access to raw radiofrequency (RF) channel data and is typically confined to high-frequency acquisitions (typically 20-40 MHz) [18]. By contrast, the sparse deconvolution approach pursued here operates directly on the SVD-filtered power-Doppler frames of the standard contrast-free pipeline: it needs neither contrast agents nor RF-channel access, and it remains robust to the residual noise that dominates label-free Doppler at depth. Notably, the same sparse-deconvolution framework has recently been carried into contrast-enhanced ultrasound: FLAME combines a sixth-order temporal cumulant of microbubble fluctuations with Richardson–Lucy and sparsity-and- continuity deconvolution to reach about 50 µm in three dimensions [19]. Its resolution gain, however, originates in the high-order statistics of injected microbubbles, whereas the framework pursued here is contrast-free and asks how far the sparsity and continuity priors alone can be taken on SVD-filtered power Doppler, extending them to velocity-resolved and task-evoked functional imaging.

In this study, we establish a sparse deconvolution framework to SVD-filtered ultrafast power Doppler imaging and assess its ability to jointly enhance contrast and resolution. The contributions of this work are fourfold. First, we transfer and adapt the sparsity-and-continuity sparse-deconvolution framework from fluorescence microscopy to contrast-free SVD power Doppler and validate it on a numerical phantom with known ground truth. Second, we systematically dissect the individual roles of the sparsity and continuity priors through a four-way ablation— conventional SVD, sparsity-only, continuity-only, and combined sparsity-with-continuity— evaluated in vivo with a panel of background-independent image-quality metrics, so as to characterize, rather than overstate, the practical gains attainable without contrast agents. Third, we introduce a velocity-band decomposition that conditions the deconvolution on flow speed, yielding a velocity-resolved microvascular map. Fourth, we extend the framework from structural to functional imaging by applying it to task-evoked functional ultrasound, and show that it preserves the stimulus-evoked haemodynamic response while sharpening the spatial localization of the activation map. Throughout, we benchmark against conventional SVD power Doppler on in vivo rat-brain plane-wave acquisitions obtained with a 15-MHz linear array on a Verasonics Vantage system.

## 2. Materials and Methods

### 2.1. Animal preparation and ultrafast ultrasound acquisition

Adult male Sprague–Dawley rats were used in compliance with the European Communities Council Directive (2010/63/EU) and the local ethical committee for animal research. Anaesthesia was induced by an intraperitoneal bolus of medetomidine (0.4 mg kg⁻¹) and ketamine (40 mg kg⁻¹), and a saline-filled catheter was inserted into the jugular vein before the animal was placed on a stereotaxic frame. A craniotomy was then performed between Bregma and Lambda to provide acoustic access, leaving the dura mater intact. Approximately 45 min after induction, once the craniotomy was completed, anaesthesia was maintained at a reduced level by subcutaneous perfusion of medetomidine (0.1 mg kg⁻¹ h⁻¹) and ketamine (12.5 mg kg⁻¹ h⁻¹) delivered with a syringe pump. Body temperature was maintained at 37 °C with a heating blanket and an intrarectal probe, and heart and respiratory rates were monitored continuously to verify the stability of the anaesthesia throughout the imaging session.

Ultrafast ultrasound acquisitions were performed with a Verasonics Vantage 256 research scanner driving a customized 15-MHz linear-array probe coupled to the imaging window with degassed ultrasound gel. The probe contained 128 elements at a pitch of 110 µm. Coherent plane-wave compounding [20] was implemented with 11 tilted plane waves spanning angles from −10° to +10° in 2° steps, transmitted at a pulse repetition frequency of 5500 Hz. Compounded frames were thus generated at an effective frame rate of 500 Hz. Multiple acquisitions were made across coronal and sagittal planes.

### 2.2. Beamforming and SVD-based clutter filtering

Radiofrequency (RF) channel data were demodulated to baseband in-phase and quadrature (IQ) signals and beamformed by a delay-and-sum (DAS) algorithm. The beamformed IQ stack was reshaped into a Casorati matrix of size (Nₓ·Nz) × Nt, where Nₓ and Nz are the spatial dimensions and Nt is the number of compounded frames in the block.

Tissue clutter, blood signal, and noise were separated by a spatiotemporal SVD of the Casorati matrix, applied independently to each block of 200 consecutive compounded frames (400 ms). Within every block the singular-value cutoff was fixed at 10% of the block length: the first 20 singular components, which carry the spatially and temporally coherent tissue clutter, were discarded, and singular components 20 to 200 were retained and summed to reconstruct a filtered IQ stack. The same 10% threshold was used for every dataset in this study, so that no acquisition-specific tuning of the clutter filter enters the comparison between reconstruction strategies. A power-Doppler frame was then obtained for each block by integrating the squared magnitude across the temporal dimension, and the full acquisition was thus represented as a stack of SVD-filtered power-Doppler frames, which served as the input to the sparse deconvolution stage.

### 2.3. Sparse deconvolution framework

We adapted the sparse deconvolution framework originally developed for fluorescence super-resolution microscopy to SVD-filtered power-Doppler images. Let f denote a single SVD-filtered Doppler frame, A is the system PSF acting as a linear blurring operator, b is a low-spatial-frequency background estimate, and x is the unknown high-resolution image. The reconstruction is formulated as the convex minimization problem:

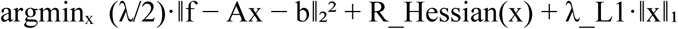

The first term enforces data fidelity between the recovered image x and the measured Doppler frame f. The second term, R_Hessian(x), is a Hessian continuity regularizer that penalizes large second-order spatial derivatives along the lateral, axial, and temporal axes, thereby suppressing noise and reconstruction artifacts while preserving smooth vessel structures. The third term is an ℓ1 sparsity prior that exploits the intrinsic sparsity of the microvascular bed at the imaging resolution and promotes the recovery of high-frequency components inside the system OTF. Two scalar weights, λ (fidelity) and λ_L1 (sparsity), balance the contributions of the three terms. As reported in [9], optimal values of these two weights follow an approximately linear relationship and depend on the SNR of the input data.

The optimization was solved by a Split-Bregman scheme, which alternates closed-form proximal updates for the data-fidelity term, the Hessian continuity term, and the soft-thresholding operator associated with the ℓ1 prior. All large-scale convolutions involved in the Hessian and PSF operators were precomputed in the Fourier domain, where the Hessian eigenvalues were stored as fixed kernels and the PSF was represented through its OTF. Convolution and adjoint operations were implemented as inline anonymous functions to avoid repeated memory allocations during the iterative loop. The background term b was estimated by a multilevel two-dimensional wavelet decomposition of the input frame, retaining only the lowest-frequency band before back-projection, as described in [9].

After the iterative reconstruction, an accelerated Richardson–Lucy deconvolution [21] with the estimated system PSF was applied to recover sub-PSF detail while enforcing non-negativity. We used the Andrews–Biggs vector-extrapolation scheme [22] to accelerate convergence and limit the number of updates required to reach a stable solution. The system PSF was estimated from a calibration acquisition of a 50-µm tungsten-wire target imaged in a water tank with the same transmit/receive sequence as the in vivo acquisition. The lateral and axial profiles of the resulting point image were fitted to a separable two-dimensional Gaussian, which served as the PSF model A in both the iterative reconstruction and the Richardson–Lucy deconvolution.

The full pipeline is summarized as follows: (i) DAS beamforming of the plane-wave-compounded RF data into IQ frames; (ii) spatiotemporal SVD clutter filtering and power-Doppler integration; (iii) wavelet-based background estimation; (iv) Split-Bregman iterative reconstruction with sparsity and Hessian continuity priors; and (v) accelerated Richardson–Lucy deconvolution with the calibrated PSF. Pixel upsampling by a factor of two was optionally applied between steps (iii) and (iv) to mitigate the effects of finite Nyquist sampling, as suggested in [9].

### 2.4. Implementation and parameter selection

The full pipeline was implemented in MATLAB R2023a (The MathWorks, Natick, MA, USA) and executed on a workstation equipped with an Intel Xeon W-2295 CPU and an NVIDIA RTX A6000 GPU. Computationally intensive steps—including the SVD of the Casorati matrix, the Hessian and PSF convolutions, and the Richardson–Lucy deconvolution—were dispatched to the GPU using the MATLAB Parallel Computing Toolbox and CUDA-accelerated FFT routines.

The fidelity weight λ and the sparsity weight λ_L1 were the only content-aware parameters that required tuning. Following the procedure recommended in [9], we adjusted them by visual inspection of the reconstruction so that small vessels became sharper without the appearance of high-frequency artifacts or the suppression of weak signals. For the in vivo rat-brain dataset reported here, we converged on a high-fidelity, low-sparsity regime, which preserved the dynamic range of the SVD-filtered input while still extracting capillary-scale features. The number of Split-Bregman iterations was fixed at 100 and the number of accelerated Richardson–Lucy iterations at 10. All other parameters (background-estimation level, Hessian regularization weight, upsampling factor) were left at the default values published with the original sparse deconvolution software [9].

### 2.5. Numerical phantom simulation

To evaluate the four reconstruction strategies against a known ground truth, we generated a two-dimensional numerical microvascular phantom on the same spatial grid and field of view as the in vivo acquisitions. The phantom contained a set of curved vessels distributed across depth, each defined as a thin tubular path with a randomly perturbed trajectory; superficial vessels were assigned higher amplitudes than deep vessels to reproduce the depth-dependent loss of blood signal. The binary union of all vessel paths was retained as the ground-truth vessel map. Each vessel was blurred by the system PSF model and attenuated with depth, and was modulated over slow time by a complex exponential with a randomly assigned Doppler frequency to mimic flowing blood. Tissue clutter was modeled as a small number of smooth, spatially low-rank patterns evolving slowly over time, and complex Gaussian noise was added, yielding a synthetic 500-frame complex dataset.

The synthetic dataset was processed by the same pipeline used for the in vivo data. Tissue clutter was removed by discarding the leading singular components of the spatiotemporal SVD, and the four reconstruction strategies—conventional SVD power Doppler, sparsity-only, continuity-only, and combined sparsity-with-continuity sparse deconvolution—were obtained by enabling or disabling the corresponding regularization terms in the Split-Bregman optimization. Because the ground-truth vessel positions were known, the simulation permitted direct measurement of vessel width and of resolution-related metrics that cannot be defined unambiguously in vivo. We note that the phantom vessels were blurred with the same Gaussian PSF model later used in the reconstruction; the simulation therefore validates the optimization and the relative ordering of the priors under matched-PSF conditions, and is complemented by the in vivo evaluation below.

### 2.6. Quantitative metrics

Unless stated otherwise, all quantitative metrics were computed on power-Doppler images normalized to their own maximum, so that the comparison reflected the spatial distribution of intensity rather than the absolute scale, which differs between methods. Let PD denote a power-Doppler image with pixel values pᵢ (i = 1 … N); its max-normalized version is

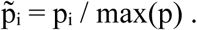

Six image-quality metrics and two frame-dependence and spectral analyses were used, defined as follows.

#### Gini coefficient (sparsity)

Sparsity was quantified by the Gini coefficient of the normalized intensities. Sorting the N pixel values in ascending order as pȃ(1) ≤ … ≤ p̂(N), the coefficient was computed as

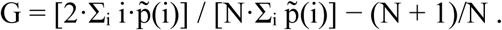

It ranges from 0 for a uniform image to a value approaching 1 when all energy is concentrated in a single pixel; higher values indicate a sparser, more vessel-concentrated image.

#### Kurtosis (sharpness)

The sharpness of the intensity distribution was measured by its kurtosis,

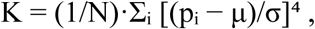

where μ and σ are the mean and standard deviation of the normalized image. Higher kurtosis reflects a more strongly peaked distribution dominated by a few bright vessel pixels over a low background.

#### Dynamic range

The dynamic range was defined as

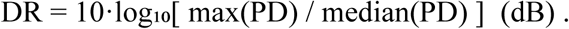

Because the median of a sparse power-Doppler image is governed by background pixels, this quantity reflects the separation between the brightest vessels and the residual background floor, with larger values indicating better background suppression relative to the vessel signal.

#### Deep-signal fraction

To quantify the preservation of deep vascular signal, the image was partitioned at an axial depth of 5 mm and the deep-signal fraction was defined as the proportion of total normalized intensity located beyond this depth,

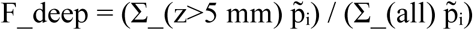

This background-independent ratio increases when more signal is recovered from deep regions relative to the whole image.

#### Vessel full-width-at-half-maximum (FWHM)

The apparent vessel width was measured as the FWHM of intensity profiles taken across vessels. For each profile the maximum was located, the half-maximum level was set to half of the peak above the local baseline, and the FWHM was obtained by linear interpolation of the two half-maximum crossings on either side of the peak; lateral and axial FWHM were measured from profiles taken along the lateral and axial directions, respectively. In the numerical phantom, profiles were centered on each ground-truth vessel and the FWHM was reported per vessel; in vivo, where the true vessel positions are unknown, the FWHM was measured on the strongest vessel within a region of interest and pooled across acquisition blocks. We treat the FWHM as a comparative apparent-width measure rather than a calibrated point-spread resolution (see Discussion). In addition, normalized lateral intensity profiles extracted at a fixed depth were used to visualize vessel separation and the residual background level.

#### Frame-number dependence (convergence)

To assess how image quality builds up with acquisition length, power-Doppler images were accumulated from increasing numbers of frames, sampled logarithmically between 1 and the full 5000 frames (the concatenation of all 25 blocks). For each method, the full 5000-frame accumulation served as that method’s own reference, and convergence was quantified by the Pearson correlation coefficient between the max-normalized N-frame image and the corresponding reference. The same frame sampling was used to track the evolution of the Gini coefficient and of the deep-signal fraction with frame number.

#### Spatial-frequency content

To compare the recovery of fine structures, the two-dimensional power spectrum of each full-accumulation power-Doppler image was computed as the squared magnitude of its discrete Fourier transform, after removal of the mean and apodization with a separable Hann window to limit spectral leakage. The frequency axes were converted to physical units (cycles·mm⁻¹) using the pixel spacing of each reconstruction. An azimuthally averaged (radial) power spectrum was then obtained by averaging the two-dimensional spectrum within concentric frequency rings; a broader radial spectrum and higher power at high spatial frequencies indicate the recovery of finer spatial detail.

### 2.7. Functional ultrasound acquisition and activation mapping

To evaluate sparse deconvolution in a functional setting, task-evoked cerebral haemodynamics were imaged in the somatosensory cortex during whisker stimulation, using the same probe, plane-wave compounding, beamforming and SVD clutter filtering as above; the acquisition followed the protocol described previously [23]. The facial whiskers contralateral to the imaged hemisphere were deflected by a custom mechanical stimulator brushing the whiskers at 10 Hz. The stimulator was driven by a servomotor under microcontroller (Arduino Uno) control and triggered by the imaging sequence, ensuring synchronisation between stimulation and image acquisition and reproducible stimulus timing across repetitions. Power-Doppler frames were acquired continuously as consecutive blocks of 200 compounded frames (400 ms per block, i.e. 2.5 power-Doppler frames per second), each block yielding one SVD-filtered power-Doppler frame. Stimulation followed a block design of five cycles of 40 s of rest followed by 30 s of stimulation (70 s per cycle); the stimulation paradigm therefore lasted 350 s and was represented as a temporal sequence of 875 power-Doppler frames. Because the power-Doppler signal is proportional to the moving-blood volume within a voxel and hence to the local cerebral blood volume (CBV) [3], the relative CBV change was computed per block as ΔCBV/CBV₀ = (PD(t) − PD₀)/PD₀ × 100, where PD₀ is the mean power-Doppler signal over the baseline (rest) periods. The identical time series was reconstructed by sparse deconvolution, so that the conventional SVD and the sparse-deconvolution functional maps could be compared on the same acquisition.

Activation maps were computed following the correlation approach of functional ultrasound [3]. For every pixel, the Pearson correlation coefficient r between its power-Doppler time course and the binary stimulus regressor A(t) (one during stimulation, zero during rest) was calculated and converted to a z-score by the Fisher transformation, z = √(Nₜ − 3)·arctanh(r), where Nₜ is the number of time points; pixels with z > 3.1 (p < 0.001) were labelled as activated and overlaid on the power-Doppler anatomy. A region of interest enclosing the activated barrel cortex was then defined, and the mean ΔCBV/CBV₀ time course within it was extracted for both reconstructions; the response was summarized as the mean ΔCBV/CBV₀ during the rest and stimulation phases. Because sparse deconvolution is a nonlinear operation, the sparse-deconvolution ΔCBV/CBV₀ magnitude is treated as a spatially resolved qualitative index rather than a calibrated haemodynamic measure, whereas the correlation-based activation maps, being invariant to monotonic intensity transformations, provide a direct comparison of spatial specificity (see Discussion).

### 2.8. Ultrasound localization microscopy

The ULM data used for the comparison in Section 3.6 were acquired in the same animal and in the same imaging plane as the contrast-free acquisition, immediately after it, and were processed as described previously [23]. SonoVue microbubbles (Bracco, reconstituted in 5 mL of saline) were infused continuously into the jugular vein at 3.5 mL h⁻¹ with a push syringe over approximately 20 min, corresponding to a total injected volume of about 1.1 mL; a magnet placed inside the syringe kept the suspension mixed throughout the acquisition. Contrast-mode data were acquired as continuous blocks of 400 compounded frames at a 1000 Hz frame rate, each compounded frame being formed from five tilted plane waves (−5°, −2°, 0°, +2° and +5°) fired at a 5000 Hz pulse repetition frequency, with a two-cycle transmit pulse and a mechanical index of 0.09.

Beamformed IQ data were filtered with the same spatiotemporal SVD clutter filter, discarding the first ten singular components, and interpolated with a Lanczos kernel onto a grid one sixth of the probe pitch laterally and one sixth of the wavelength axially. Microbubbles were detected as local intensity maxima whose neighbourhood correlated at more than 0.7 with a Gaussian model of the system point spread function, and their positions were refined to sub-pixel accuracy by a second-order polynomial fit over a 5 × 5 pixel neighbourhood and rounded onto a 6.875 × 6.25 µm grid. Detections were linked across frames by a Hungarian-assignment particle tracker without gap filling, the maximum linking distance being set by a 100 mm s⁻¹ upper bound on microbubble speed; only tracks spanning at least ten successive frames were retained, and each track was linearly interpolated so that every pixel along the microbubble path received one count. Slow drift over the acquisition was corrected by intensity-based translational registration of microbubble- count maps computed over 10 s intervals. The structural ULM image used here is the accumulated microbubble-count map, restricted to pixels accumulating at least five independent detections.

Functional ULM maps were obtained by accumulating microbubble detections in a 5 s sliding window with a 1 s step, summing equivalent time points across the repeated stimulation cycles to form a pattern-averaged microbubble-flux sequence, and computing for each pixel the Pearson correlation between that sequence and the binary stimulus regressor, exactly as for the power-Doppler activation maps described above. Because the ULM and functional ULM images derive from a separate contrast-enhanced acquisition and are not pixel-registered to the contrast-free maps, they are used throughout as a qualitative super-resolution reference rather than as a ground truth.

### 2.9. Statistical analysis

Quantitative metrics are reported as mean ± standard error of the mean (SEM). To obtain repeated measurements, FWHM values were sampled per vessel (numerical phantom) or per acquisition block (in vivo, n = 25 blocks per method), while the Gini coefficient, kurtosis, dynamic range, and deep-signal fraction were sampled per acquisition block or, for the phantom, per depth sub-region. For each metric, an omnibus Friedman test across the four reconstruction strategies—which share the same underlying data and therefore constitute paired samples—was performed first, followed by pairwise Wilcoxon signed-rank tests with Holm correction for multiple comparisons. Differences were considered significant at p < 0.05, with significance levels denoted *p < 0.05, **p < 0.01, and ***p < 0.001.

## 3. Results

### 3.1. Simulation validation with known ground truth

To validate the four reconstruction strategies against a known ground truth, we generated a numerical microvascular phantom containing curved vessels distributed across depth, embedded in low-rank tissue clutter and additive noise (Fig. 2(a)). The simulated 500-frame dataset was processed with conventional spatiotemporal SVD filtering (Conventional SVD) and with sparse deconvolution (SD) using sparsity-only, continuity-only, or combined sparsity-with-continuity regularization (Fig. 2(b–e)). Quantitative metrics were computed per vessel (lateral and axial FWHM) and per depth block (Gini coefficient and kurtosis); group comparisons used pairwise Wilcoxon signed-rank tests with Holm correction (Fig. 2(f–i); mean ± SEM).

**Figure 1.**
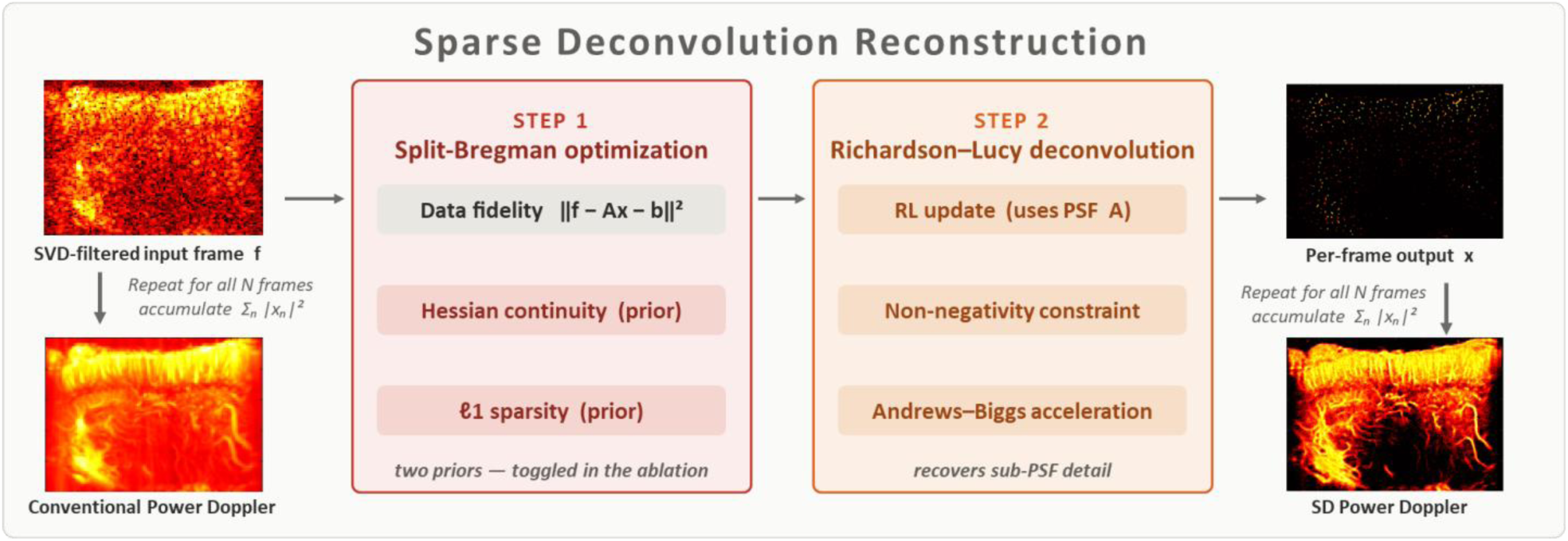
Processing pipeline of the sparse deconvolution reconstruction.

**Figure 2.**
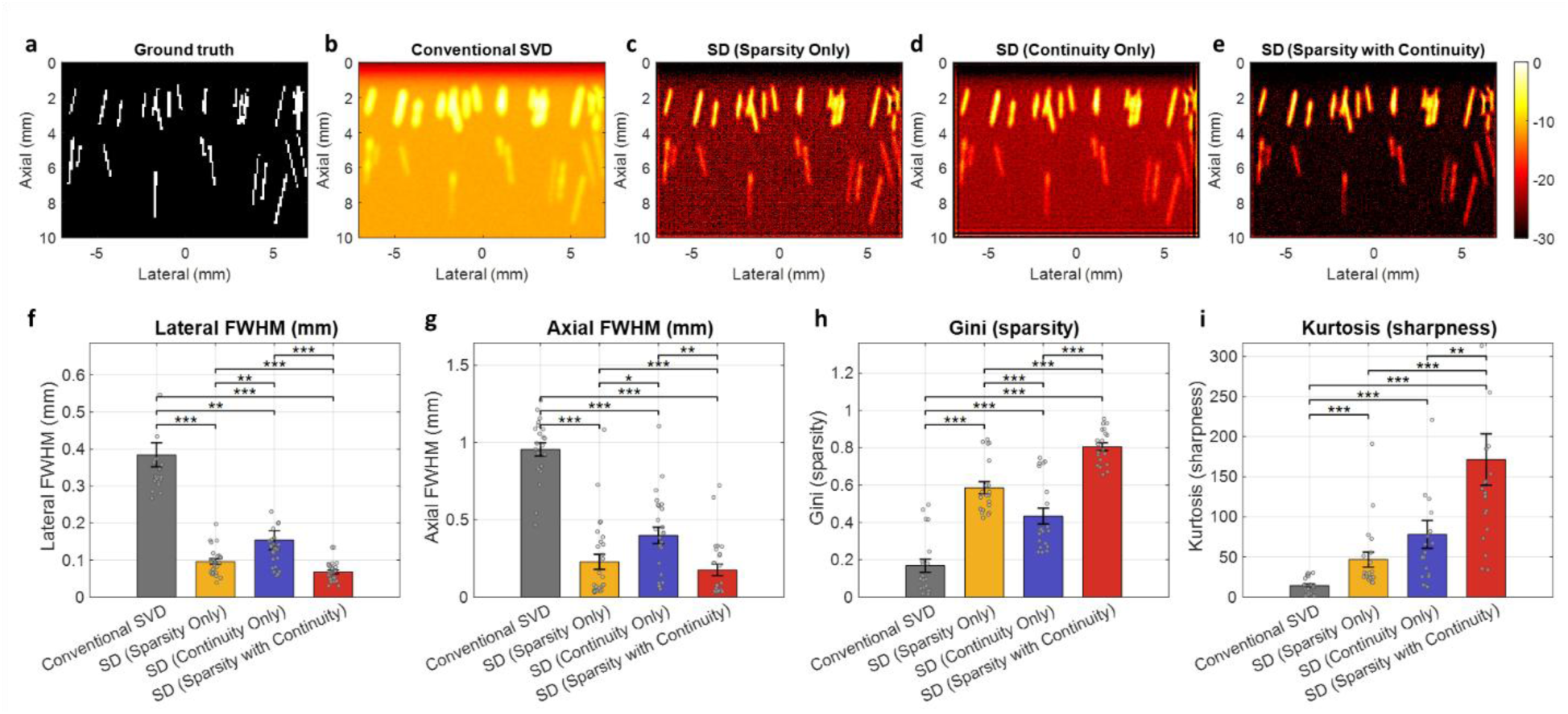
Simulation of the four reconstruction strategies against a known ground truth. (a) Ground-truth microvascular phantom (binary vessel map). (b–e) Power-Doppler reconstructions of the simulated 500-frame dataset (displayed in dB) obtained with (b) Conventional SVD, (c) SD with sparsity-only regularization, (d) SD with continuity-only regularization, and (e) SD with combined sparsity-with-continuity regularization. (f–i) Quantitative comparison: (f) lateral FWHM and (g) axial FWHM measured at each ground-truth vessel; (h) Gini coefficient and (i) kurtosis computed per depth block. Bars show mean ± SEM with individual measurements overlaid; statistical comparisons used pairwise Wilcoxon signed-rank tests with Holm correction (*p < 0.05, **p < 0.01, ***p < 0.001).

Conventional SVD produced a blurred reconstruction with strong residual background and poorly defined vessels (Fig. 2(b)), whereas all three sparse-deconvolution variants markedly sharpened the vasculature and suppressed the background (Fig. 2(c–e)). These visual differences were confirmed quantitatively. The lateral FWHM decreased from 0.384 mm (Conventional SVD) to 0.096 mm (sparsity-only), 0.154 mm (continuity-only), and 0.068 mm for the combined formulation, and the axial FWHM decreased from 0.956 mm to 0.228, 0.399, and 0.176 mm, respectively (Fig. 2(f,g); all sparse-deconvolution variants vs Conventional SVD, p < 0.001). The combined sparsity-with-continuity formulation achieved the narrowest vessels in both directions, a 5.6-fold lateral and 5.4-fold axial reduction in apparent width over Conventional SVD, and significantly outperformed both single-prior variants (p < 0.001 vs continuity-only; p < 0.05 vs sparsity-only for axial FWHM).

The sparsity and sharpness metrics showed the same ordering. The Gini coefficient increased from 0.168 (Conventional SVD) to 0.586 (sparsity-only), 0.434 (continuity-only), and 0.806 for the combined formulation, and the kurtosis increased from 14.2 to 46.7, 78.1, and 171.4, respectively (Fig. 2(h,i); all p < 0.001 vs Conventional SVD). The combined formulation again yielded the highest values, significantly exceeding both single-prior variants (p < 0.001 for Gini; p < 0.01 for kurtosis vs continuity-only). Across all four metrics, sparsity-only and continuity-only each captured part of the benefit—sparsity-only favoring resolution and continuity-only favoring sharpness of connected structures—whereas only their combination consistently ranked best, confirming on ground-truth data that the two regularization terms are complementary and that the combined sparse-deconvolution formulation provides the best overall reconstruction.

### 3.2. In vivo characterization of sparse deconvolution versus conventional SVD

We characterized sparse deconvolution against conventional spatiotemporal SVD on an in vivo rat-brain dataset (5000 compounded frames) along several complementary axes (Fig. 3). Power- Doppler images reconstructed from increasing numbers of frames (1, 10, 100, 500, and 5000) are shown for conventional SVD (Fig. 3(a)) and sparse deconvolution (Fig. 3(b)). Conventional SVD produced a continuous but blurred vascular map dominated by a diffuse background, whereas sparse deconvolution resolved individual cortical penetrating vessels and deep microvascular branches with markedly higher contrast and sharpness.

**Figure 3.**
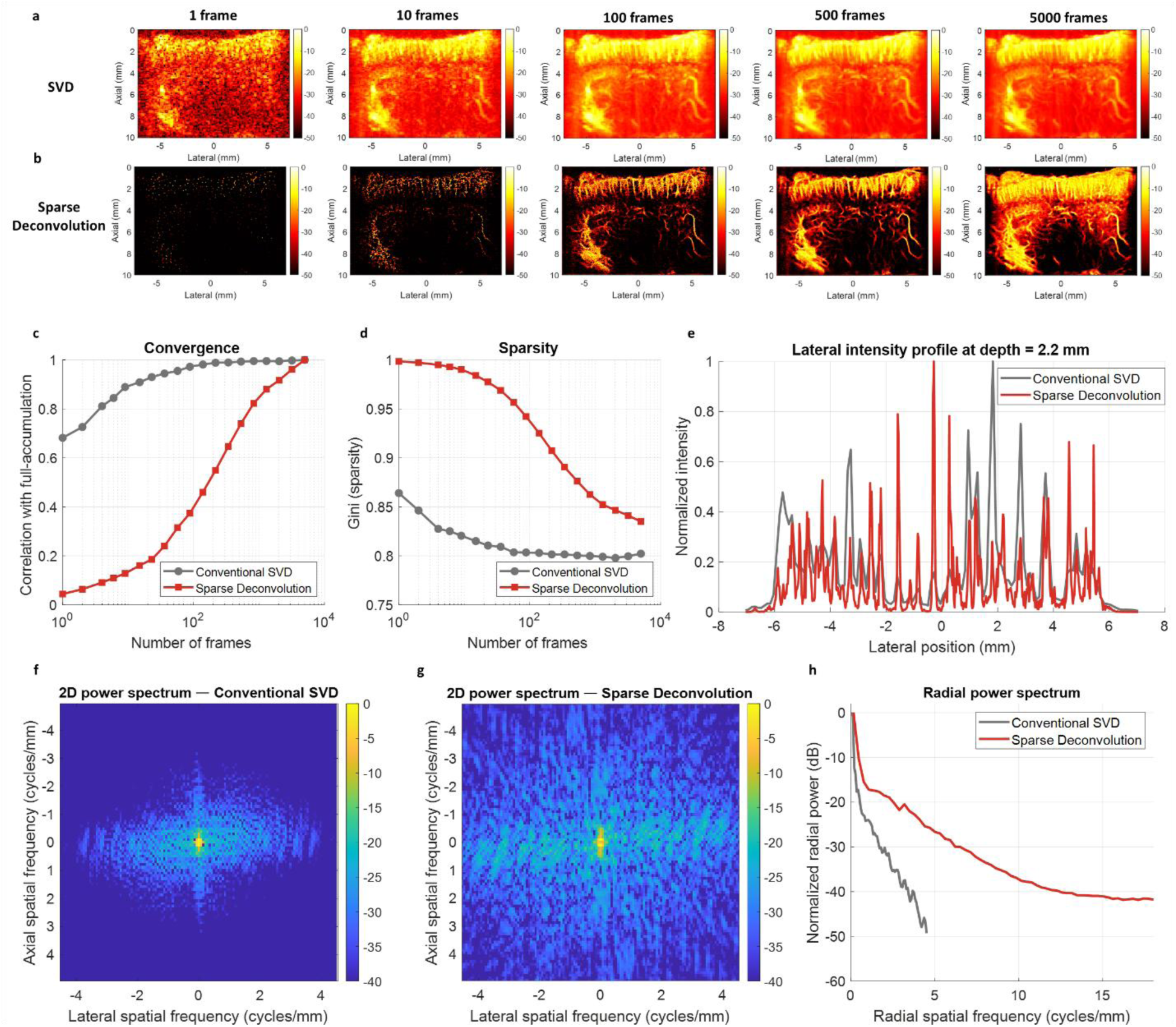
In vivo comparison of conventional SVD and sparse deconvolution. (a, b) Power-Doppler images of the rat brain reconstructed from increasing numbers of frames (1, 10, 100, 500, 5000; displayed in dB) using (a) conventional spatiotemporal SVD and (b) sparse deconvolution. (c) Correlation of each frame-limited reconstruction with its own full 5000-frame accumulation, as a function of frame number (log scale). (d) Gini coefficient (sparsity) versus frame number. (e) Normalized lateral intensity profiles extracted at a depth of 2.2 mm, illustrating vessel separation and background suppression. (f, g) Two-dimensional power spectra of the full-accumulation power-Doppler images for (f) conventional SVD and (g) sparse deconvolution, displayed on identical spatial-frequency axes (dB). (h) Radial (azimuthally averaged) power spectra of the two reconstructions. In (c, d, h), grey denotes conventional SVD and red denotes sparse deconvolution.

The two methods exhibited distinct accumulation behavior with frame number (Fig. 3(c)). Taking each method’s full 5000-frame reconstruction as its own reference, conventional SVD converged rapidly, reaching a correlation above 0.9 within ≈10 frames, whereas sparse deconvolution converged more gradually and required substantially more frames to approach its reference. This difference is consistent with the higher sparsity of the deconvolved images: because each frame contributes a small number of highly localized detections, more frames are required to progressively populate the microvascular map. The sparsity itself, quantified by the Gini coefficient (Fig. 3(d)), was consistently higher for sparse deconvolution than for conventional SVD across all frame numbers, confirming a more concentrated, vessel-specific intensity distribution.

The resolution gain was directly visible in the lateral intensity profiles extracted at a representative cortical depth (Fig. 3(e)). Sparse deconvolution produced sharp, well-separated peaks with near-zero inter-vessel valleys, whereas conventional SVD yielded broad, overlapping peaks sitting on an elevated background, indicating that closely spaced vessels unresolved by conventional SVD were separated after deconvolution. This was corroborated in the spatial-frequency domain: the two-dimensional power spectra (Fig. 3(f,g)) show that conventional SVD concentrated its energy near the spatial-frequency origin, whereas sparse deconvolution distributed energy over a substantially broader frequency support. The radial power spectrum (Fig. 3(h)) quantifies this difference, with sparse deconvolution exhibiting consistently higher power at high spatial frequencies and extending its frequency content well beyond that of conventional SVD, consistent with the recovery of fine, sub-resolution microvascular structures.

Together, these analyses show that sparse deconvolution enhances spatial resolution, sparsity, and high-frequency content relative to conventional SVD, at the cost of requiring a larger number of frames to fully populate its sparser, more detailed vascular representation.

### 3.3. Sparse deconvolution improves microvascular power-Doppler imaging in vivo

We compared four reconstruction strategies on the same in vivo rat-brain dataset: conventional spatiotemporal SVD filtering (Conventional SVD), and sparse deconvolution (SD) using sparsity regularization only, continuity regularization only, or the combined sparsity-with-continuity formulation. Representative power-Doppler maps are shown in Fig. 4(a), with magnified cortical views in Fig. 4(b). Quantitative metrics were computed per acquisition block (n = 25 blocks per method); group comparisons used the Friedman test followed by pairwise Wilcoxon signed-rank tests with Holm correction (Fig. 4(c–h)). Data are reported as mean ± SEM.

**Figure 4.**
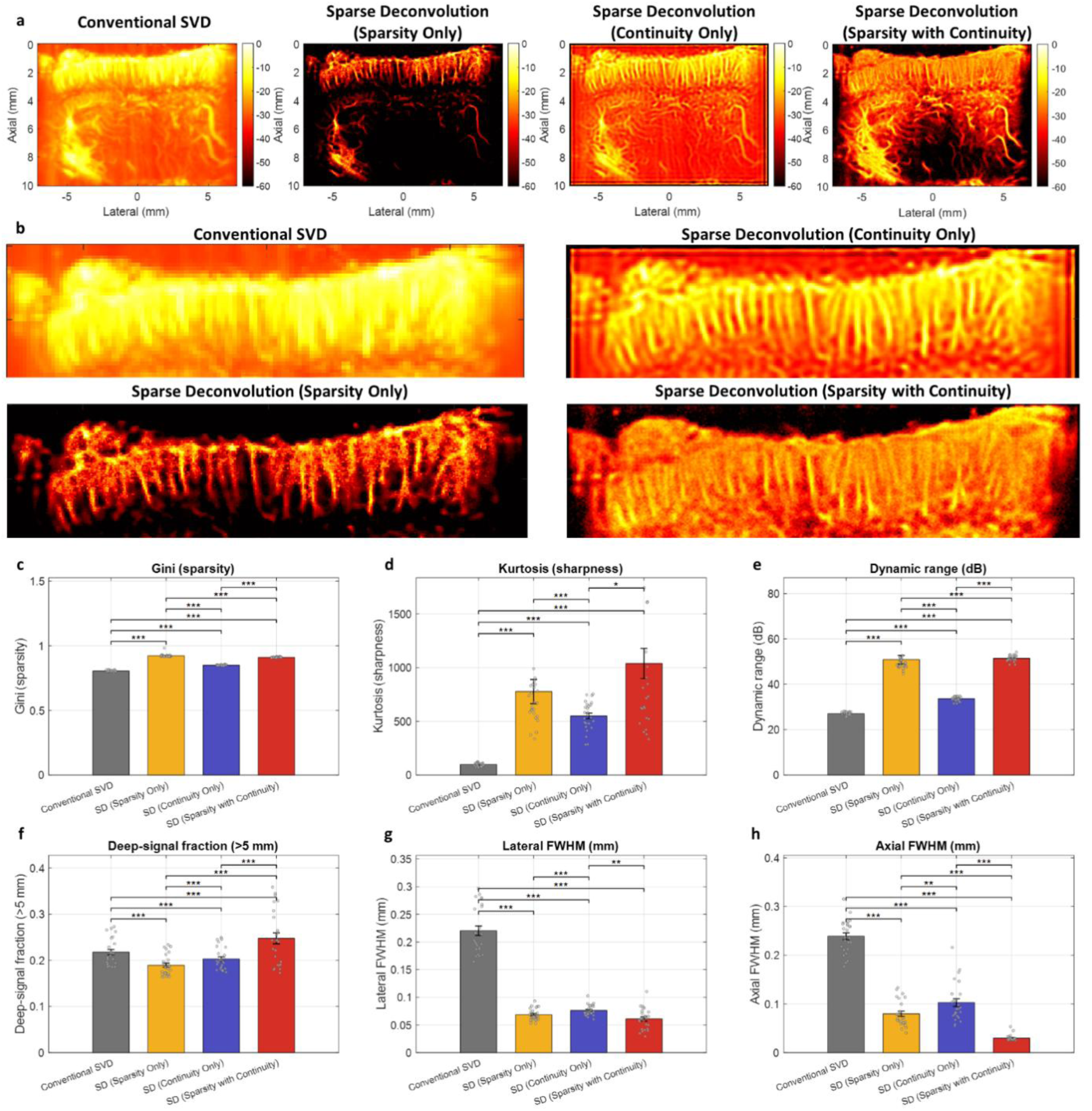
Sparse deconvolution improves in vivo microvascular power-Doppler imaging. (a) Representative power- Doppler maps (displayed in dB) reconstructed with conventional spatiotemporal SVD and with sparse deconvolution using sparsity-only, continuity-only, and combined sparsity-with-continuity regularization. (b) Magnified cortical views of the same reconstructions. (c–h) Per-block quantitative comparison (n = 25 blocks per method; mean ± SEM; individual blocks overlaid as points): (c) Gini coefficient (sparsity), (d) kurtosis (sharpness), (e) dynamic range, (f) deep-signal fraction beyond 5 mm depth, (g) lateral FWHM, and (h) axial FWHM. Statistical comparisons used the Friedman test with pairwise Wilcoxon signed-rank tests and Holm correction (*p < 0.05, **p < 0.01, ***p < 0.001).

Compared with Conventional SVD, all three sparse-deconvolution variants markedly increased the sparsity and sharpness of the reconstructed vasculature. The Gini coefficient increased from 0.805 ± 0.001 (Conventional SVD) to 0.911 ± 0.001 for the combined SD (Fig. 4(c)), and the image kurtosis increased from 97 ± 3 to 1038 ± 140 (Fig. 4(d)), indicating a strongly peaked, vessel-concentrated intensity distribution. The dynamic range rose correspondingly from 27.1 ± 0.1 dB to 51.5 ± 0.3 dB (Fig. 4(e)). These improvements were highly significant for every sparse-deconvolution variant relative to Conventional SVD (all p < 0.001).

The apparent vessel width narrowed in parallel. The lateral FWHM of the smallest resolved vessels decreased from 0.220 ± 0.009 mm (Conventional SVD) to 0.061 ± 0.004 mm for the combined SD (Fig. 4(g)), and the axial FWHM decreased from 0.239 ± 0.007 mm to 0.030 ± 0.001 mm (Fig. 4(h)), a 3.6-fold and 8.0-fold reduction, respectively (both p < 0.001). Notably, the combined sparsity-with-continuity formulation achieved the narrowest axial FWHM of all methods, significantly outperforming both sparsity-only (0.080 ± 0.005 mm, p < 0.001) and continuity-only (0.103 ± 0.008 mm, p < 0.001) regularization. We emphasize that these FWHM values quantify the apparent width of the reconstructed vessels rather than a calibrated point-spread resolution; the smallest values fall below the diffraction-limited resolution of the 15-MHz array (axial wavelength ≈ 0.1 mm) and therefore reflect the intensity concentration imposed by the sparsity-promoting reconstruction rather than genuine recovery of sub-wavelength structure. They should accordingly be interpreted as comparative sharpness measures across methods on identical data.

Critically, these gains were achieved without sacrificing deep vascular signal. The fraction of signal recovered beyond 5 mm in depth was highest for the combined SD (0.248 ± 0.012), significantly exceeding Conventional SVD (0.218 ± 0.006, p < 0.001), sparsity-only (0.189 ± 0.005, p < 0.001), and continuity-only (0.203 ± 0.005, p < 0.001) reconstructions (Fig. 4(f)). In contrast, sparsity-only regularization, while reaching the highest Gini coefficient (0.923 ± 0.003), exhibited the lowest deep-signal fraction, indicating that aggressive sparsity promotion alone suppresses weak deep microvessels. The continuity term mitigated this loss: adding continuity to the sparsity prior recovered the deep-signal fraction while further sharpening vessels, as visually confirmed by the more complete deep cortical and subcortical vascular trees in Fig. 4(a, b).

Taken together, the combined sparsity-with-continuity sparse deconvolution provided the best overall balance, simultaneously achieving the highest dynamic range, the finest axial apparent width, and the greatest deep-signal preservation among all tested methods, whereas single-prior variants improved sharpness at the expense of either deep-signal sensitivity (sparsity-only) or dynamic range and sharpness (continuity-only).

### 3.4. Velocity-resolved sparse deconvolution of contrast-free cerebral blood flow

SVD-filtered power Doppler of the rat cortex (Fig. 5(a)) superimposes vascular signals spanning a wide range of flow speeds, which weakens the sparsity assumption underlying the deconvolution. To separate these populations we transformed the spatiotemporal stack into the joint spatial-temporal-frequency domain, in which a scatterer moving at speed v concentrates its energy along the cone f_t = v·|k_spatial|. The (k_z, f_t) power spectrum averaged over k_x (Fig. 5(b)), overlaid with iso-velocity lines from 1 to 150 mm/s, shows that signal energy fans out across a broad continuum of velocities rather than collapsing onto a single line, confirming the mixed-velocity composition of the cortical microvasculature.

**Figure 5.**
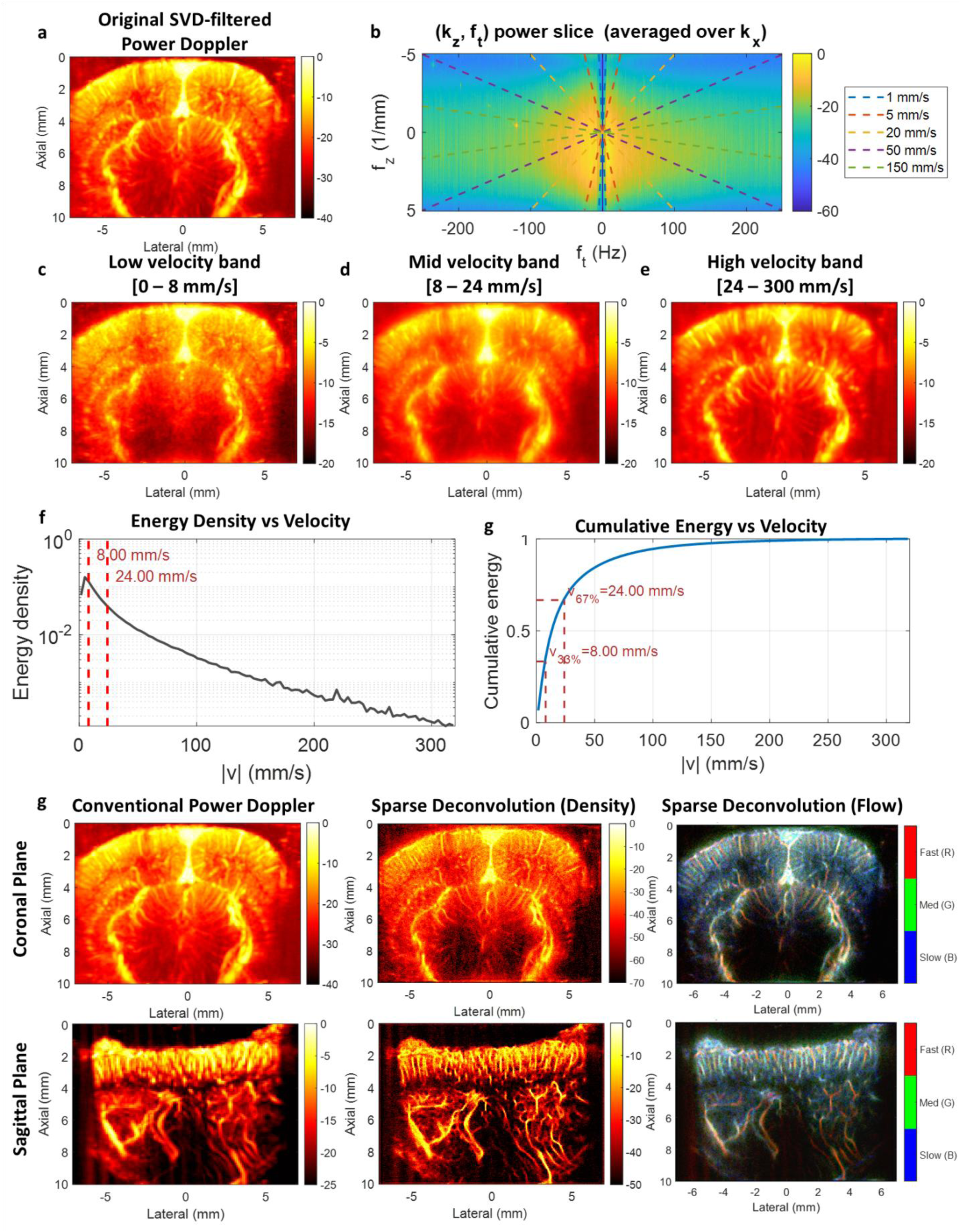
Velocity-band decomposition and velocity-resolved sparse deconvolution of contrast-free functional ultrasound in the rat brain. (a) SVD-filtered power Doppler of the coronal cortex (log scale, [−40, 0] dB). (b) Power spectrum in the (k_z, f_t) plane, averaged over the lateral spatial frequency k_x ([−60, 0] dB); dashed lines mark constant apparent speeds (1, 5, 20, 50, 150 mm/s) along which moving scatterers deposit their energy. (c–e) Power- Doppler images of the three velocity bands obtained by 3D-FFT fan filtering—low (0–8 mm/s), mid (8–24 mm/s), and high (24–300 mm/s)—each displayed over [−20, 0] dB. (f) Power-weighted velocity histogram (log ordinate); red dashed lines indicate the band edges at 8.0 and 24.0 mm/s. (g) Cumulative Doppler energy versus velocity; the 33% and 67% equal-energy tertiles define the 8.0 and 24.0 mm/s edges used in (c–e). (h) Comparison of conventional power Doppler (left), sparse-deconvolution density map (middle), and the velocity-encoded RGB composite (right; R = fast, G = medium, B = slow) for the coronal (top) and sagittal (bottom) planes. Axes are in physical units (mm); field of view 10 mm (axial) × 14.08 mm (lateral).

The power-weighted velocity histogram (Fig. 5(f)) peaks at low speeds and decays monotonically toward higher velocities. Using the cumulative-energy curve (Fig. 5(g)), we partitioned the spectrum into three equal-energy bands at the 33% and 67% tertiles, giving band edges at 8.0 and 24.0 mm/s, so that each band carries approximately one third of the total Doppler energy. Applying a 3D-FFT fan filter with these edges decomposed the data into low- (0–8 mm/s), mid- (8–24 mm/s), and high-velocity (24–300 mm/s) bands (Fig. 5(c–e)). The decomposition isolates distinct vascular components—with slower flow predominating in smaller cortical and peripheral vessels and faster flow concentrated in the larger penetrating conduits—and, critically, reduces the number of concurrently active vessels within each band, improving the conditioning of the subsequent per-band sparse deconvolution.

Reconstructing each band by sparse deconvolution and recombining them yielded substantially sharper vascular maps than conventional power Doppler in both coronal and sagittal planes (Fig. 5(h), columns 1 vs 2), resolving fine penetrating vessels that were blurred in the conventional image. Encoding the three deconvolved bands into the red, green, and blue channels (fast, medium, slow) produced a velocity-resolved composite (Fig. 5(h), column 3) in which vessels of differing flow regimes are spatially disentangled, providing a single map that conveys both microvascular structure and the local flow-speed regime.

### 3.5. Functional ultrasound on sparse deconvolution

Having established sparse deconvolution as a tool for structural microvascular imaging, we next asked whether the same framework benefits task-evoked functional ultrasound, in which neuronal activity is inferred from stimulus-locked changes in cerebral blood volume encoded in the power- Doppler signal. We first verified that the structural gains observed on resting acquisitions carried over to the functional recording. On the block-averaged power-Doppler image acquired during whisker stimulation, sparse deconvolution reproduced the cortical and subcortical vascular architecture of the conventional SVD image while markedly sharpening individual penetrating vessels (Fig. 6(a)). This sharpening was confirmed in the spatial-frequency domain: the radially averaged spatial-frequency spectrum of the sparse-deconvolution image extended to substantially higher spatial frequencies than that of the conventional image (Fig. 6(b)), indicating recovery of finer spatial detail and mirroring the resolution improvement quantified on the structural data (Section 3.3).

**Figure 6.**
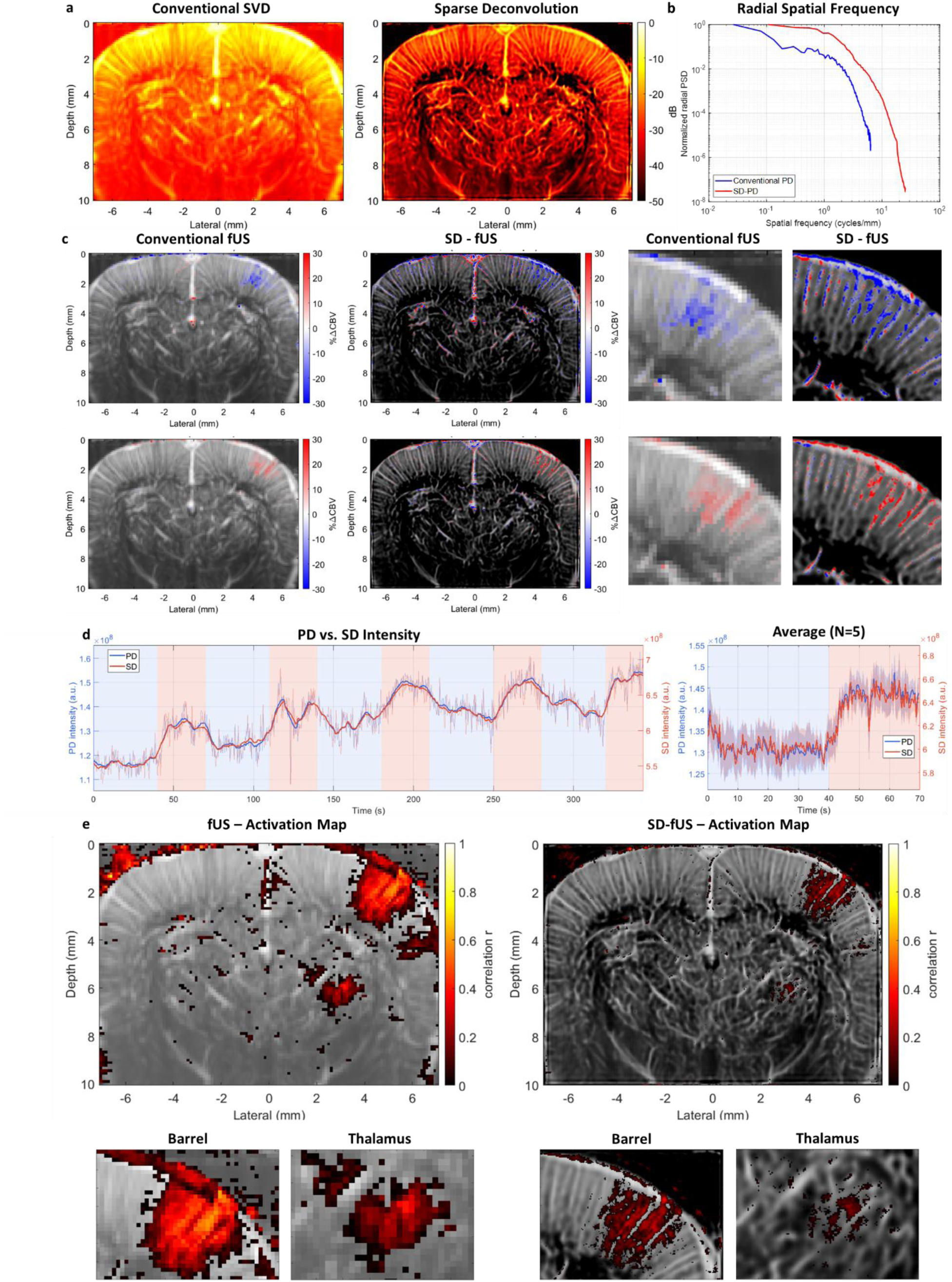
Functional ultrasound mapping on sparse-deconvolution power Doppler. Task-evoked cerebral haemodynamics were imaged in the somatosensory cortex during whisker stimulation (block design, five cycles of 40 s rest and 30 s stimulation). (a) Block-averaged structural power-Doppler images reconstructed by conventional SVD (left) and by sparse deconvolution (right), displayed in dB over a [−40, 0] dB dynamic range. (b) Radially averaged spatial-frequency spectra (normalized power spectral density, dB) of the two images in (a); the sparse-deconvolution spectrum extends to higher spatial frequencies, indicating recovery of finer spatial detail. (c) Spatial maps of the relative cerebral-blood-volume change (ΔCBV/CBV₀, %; colour scale ±30%) for conventional functional ultrasound (fUS) and sparse-deconvolution functional ultrasound (SD-fUS), overlaid on the grey-scale vascular anatomy (two representative stimulation cycles, top and bottom rows), with magnified views of the activated barrel-cortex region in the two rightmost columns. (d) Mean ΔCBV/CBV₀ time courses within the activated barrel-cortex region of interest for PD and SD across the five stimulation cycles (left; shaded bands mark the stimulation epochs) and the corresponding cycle-averaged response (right; N = 5 cycles). (e) Correlation-based functional activation maps (Pearson correlation coefficient r between each pixel time course and the stimulus regressor; threshold z > 3.1, p < 0.001) for PD (left) and SD (right), overlaid on the grey-scale anatomy, with magnified insets of the barrel-cortex and thalamic activation. Image axes are in physical units (mm).

Crucially, this spatial sharpening did not distort the underlying haemodynamic signal. Within a region of interest enclosing the contralateral barrel cortex, both the conventional SVD power Doppler and its sparse-deconvolution reconstruction exhibited a clear stimulus-locked increase or decrease in ΔCBV, rising at the onset of each stimulation epoch and returning towards baseline during the intervening rest periods (Fig. 6(c)). The two region-of-interest time courses tracked the stimulation paradigm in close temporal agreement across all five cycles, and their cycle-averaged responses were closely matched in onset latency and temporal profile (Fig. 6(d)). The relative CBV response measured after sparse deconvolution was therefore consistent, cycle to cycle, with that measured by conventional functional ultrasound, confirming that the reconstruction preserves the temporal haemodynamics of the signal rather than introducing spurious dynamics. Because the deconvolution is nonlinear, the absolute ΔCBV/CBV₀ amplitude of the sparse-deconvolution reconstruction is interpreted as a spatially resolved qualitative index rather than a calibrated haemodynamic measure (Section 2.7), whereas its temporal profile provides a faithful readout of the response dynamics.

The spatial structure of the activation map, by contrast, differed markedly between the two reconstructions. The conventional correlation map (Pearson correlation with the binary stimulus regressor, thresholded at 0.001) delineated a spatially diffuse activation cluster spread across the somatosensory cortical grey matter (Fig. 6(e)), consistent with the point-spread-limited resolution of conventional functional ultrasound, in which the mapped response reflects the blurred envelope of many neighbouring vessels rather than the vessels themselves. Applying the identical correlation analysis to the sparse-deconvolution time series confined the same response to discrete penetrating vessels within the activated territory, substantially reducing the apparent activated area while preserving its cortical location, its laminar extent, and its peak correlation value.

This gain in specificity was evident at both cortical and subcortical depths. Magnified views of the activated barrel cortex showed that the diffuse cortical patch of the conventional map resolved, after sparse deconvolution, into a set of radially oriented penetrating vessels consistent with the columnar vascular organization of the barrel field (Fig. 6(e)). A comparable sharpening was observed for the deeper thalamic activation, where the deconvolution localized the task-evoked signal to individual thalamocortical feeding vessels that were unresolved in the conventional map. Importantly, this deep response was retained rather than suppressed by the reconstruction, in line with the deep-signal preservation quantified on the structural data (Section 3.3) and attributable to the Hessian continuity prior.

Taken together, these results show that sparse deconvolution transfers its structural benefit to the functional domain. It preserves the stimulus-locked cerebral-blood-volume response measured by conventional functional ultrasound while transforming the corresponding activation map from a diffuse cortical region into a vessel-resolved map of the task-evoked haemodynamic response, and it does so entirely as a post-processing step, without any contrast agent or modification of the acquisition sequence. We further show in the Supplementary Material that the same task-related component can be recovered without any prior knowledge of the stimulus timing, directly from a singular value decomposition of the power-Doppler time series (Supplementary Section S1, Fig. S1).

### 3.6. Comparison with Ultrasound Localization Microscopy

Finally, we placed sparse deconvolution on the same footing as ultrasound localization microscopy (ULM), the reference method for super-resolved microvascular imaging, by comparing conventional power Doppler, sparse-deconvolution-enhanced power Doppler, and ULM on a matched coronal plane of the rat brain, both structurally and functionally (Fig. 7). ULM was reconstructed from a separate microbubble-contrast acquisition on the same imaging plane and serves here as a qualitative super-resolution reference for the appearance of the microvascular bed and of its functional activation, rather than as a pixel-registered ground truth for the contrast-free reconstructions.

**Figure 7.**
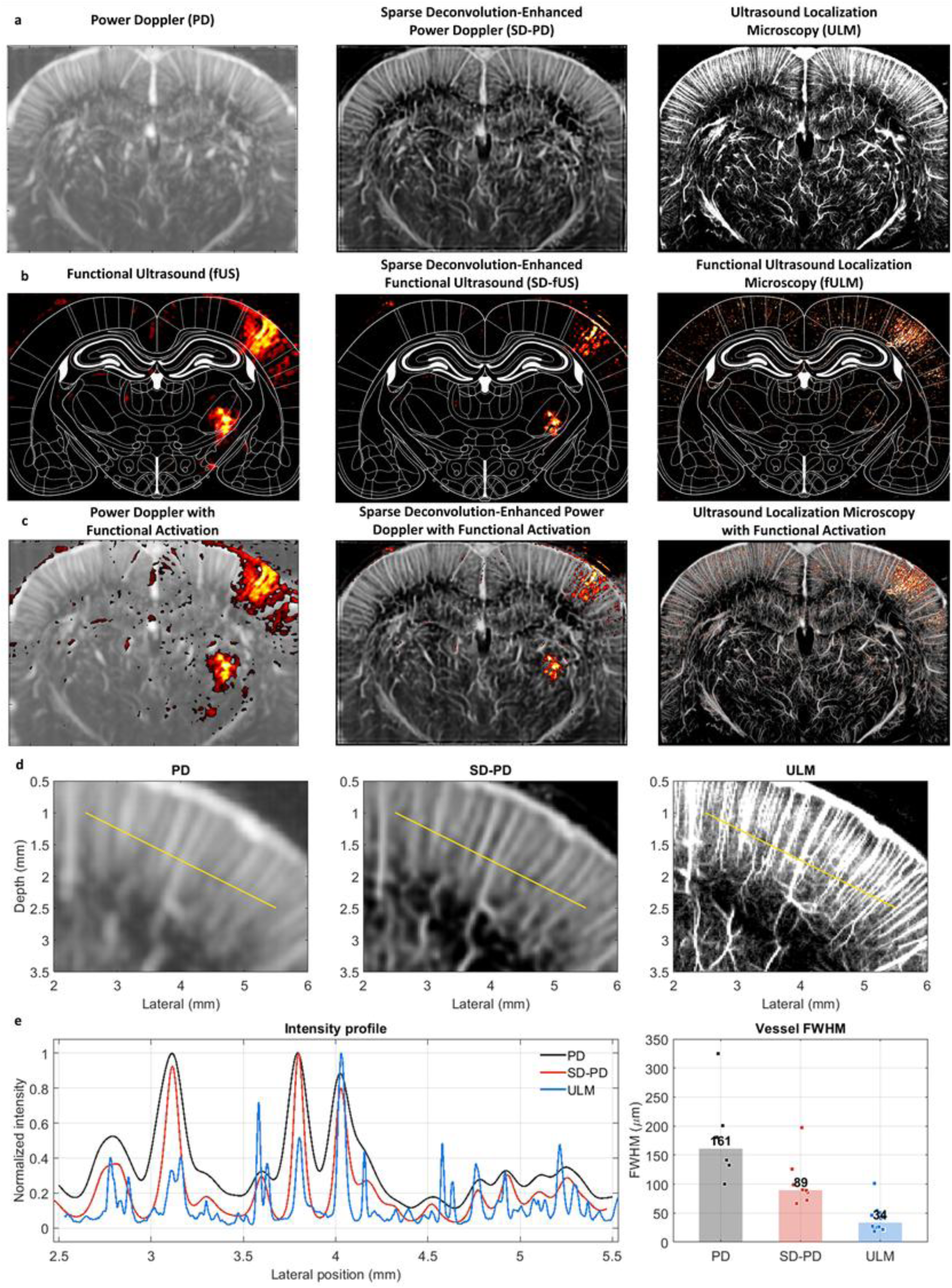
Comparison of conventional power Doppler (PD), sparse-deconvolution-enhanced power Doppler (SD- PD), and ultrasound localization microscopy (ULM) on a matched coronal plane of the rat brain, with their corresponding functional activation. ULM was obtained from a separate microbubble-contrast acquisition and serves as a qualitative super-resolution reference rather than a pixel-registered ground truth. (a) Structural microvascular images: PD (left), SD-PD (middle), and ULM (right), showing the resolution hierarchy from the point-spread-limited PD map, through the vessel-sharpened contrast-free SD-PD map, to the single-microbubble ULM map. (b) Corresponding functional activation maps: conventional functional ultrasound (fUS), SD-enhanced functional ultrasound (SD-fUS), and functional ULM (fULM); the contrast-free SD-fUS map recovers much of the discrete- vessel specificity of fULM, whereas conventional fUS reports a diffuse territory. (c) Functional activation overlaid on the respective structural image for each method, showing co-localization of the task-evoked signal with identifiable penetrating vessels for SD and ULM. (d) Zoomed-in sections of PD (left), SD-PD (middle), and ULM (right). (e) Intensity profiles across the same region of interest for the comparison of image resolution and the corresponding FWHM quantification.

Structurally, the three methods formed a clear resolution hierarchy (Fig. 7(a)). Conventional power Doppler depicted the vascular bed as a smooth, point-spread-limited intensity map in which individual microvessels were merged. ULM resolved the same bed into a dense network of individually separated vessels at the finest spatial scale, revealing microvascular detail well below the diffraction limit. Sparse deconvolution occupied an intermediate position: operating on the same SVD-filtered data as conventional power Doppler and without any contrast agent, it recovered much of the vessel-level detail present in the ULM map — sharpening penetrating vessels and separating adjacent conduits — while not attaining the single-microbubble resolution of ULM. Sparse deconvolution thus approaches ULM-like structural detail at a small fraction of its acquisition and processing cost and without microbubble injection.

The same ordering held for the functional maps (Fig. 7(b)). Functional ULM, which localizes the task-evoked haemodynamic response to individual microbubble-tracked vessels, resolved activation to discrete penetrating and thalamocortical vessels. The conventional power-Doppler activation map, by contrast, reported the response as a diffuse cortical and subcortical territory. The sparse-deconvolution activation map recovered much of the single-vessel specificity of functional ULM, confining the task-evoked signal to discrete vessels within the same activated region, despite being computed entirely from contrast-free data. Sparse deconvolution therefore transfers a large part of the functional-localization benefit of ULM to the contrast-free setting, with the distinction that its intensity is a qualitative index whereas ULM additionally yields quantitative, vessel-wise flow measurements.

Overlaying the activation maps on their respective structural images confirmed that, for both sparse deconvolution and ULM, the task-evoked signal co-localized with identifiable penetrating vessels rather than with the interstitial parenchyma (Fig. 7(c)), whereas the conventional overlay attributed the response to a broad vascular territory. Together, these comparisons position sparse deconvolution as a practical, contrast-free intermediate between conventional power Doppler and ULM: it delivers much of the structural and functional vessel-level specificity of localization microscopy while retaining the speed, simplicity, and contrast-free operation of conventional ultrafast Doppler, making it well suited to settings in which ULM cannot readily be deployed.

## 4. Discussion

In this work, we adapted the sparse deconvolution framework of Zhao et al. [9] to SVD-filtered ultrafast power Doppler ultrasound and demonstrated that it jointly improves vessel-to-background contrast and apparent spatial resolution in contrast-free in vivo brain imaging. The approach is purely computational, requires no modification of the imaging hardware or acquisition sequence, and operates on the same SVD-filtered power-Doppler frames already produced by the standard ultrafast Doppler pipeline. As such, it provides an immediately deployable front-end for high-resolution microvascular imaging in any laboratory or clinical setting where ultrafast Doppler is already available.

A central observation is that the two physical priors that underpin sparse deconvolution in fluorescence microscopy, namely sparsity and continuity, also hold for SVD-filtered Doppler images. Sparsity is justified by the fact that the microvascular bed occupies only a small fraction of the imaging volume at the resolution of a 15-MHz linear array. Continuity is justified by the smooth, tubular geometry of vessels and by the spatial coherence of blood flow within them. Because these priors are content-agnostic, the same parameter regime applied successfully to superficial cortical vessels and to deep thalamic vessels, despite the very different SNR conditions of the two regions. This is consistent with the cross-modality generalization observed in optics and supports the view that sparse deconvolution captures generic features of high-resolution biomedical imaging rather than modality-specific assumptions. These two priors play complementary and mutually stabilizing roles. The sparsity prior recovers high-frequency content beyond the diffraction limit, but on its own it tends to fragment continuous vessels and to introduce spurious, point-like artifacts. The Hessian continuity prior suppresses such artifacts and confers robustness to noise, but on its own it over-smooths the reconstruction and blurs fine structure. Only when the two priors are combined is the underlying vascular structure recovered faithfully—an interplay that motivates the systematic ablation reported below.

The gain in dynamic range, which exceeded 20 dB in vivo (increasing from approximately 27 dB to 51 dB; Fig. 4(e)), is particularly relevant for contrast-free imaging. In the absence of microbubbles, sensitivity at depth is the primary bottleneck, since blood signals are intrinsically weak and tissue clutter and electronic noise dominate the unfiltered data. Conventional SVD already removes a large fraction of this clutter, but the residual noise floor remains the limiting factor for deep microvascular imaging. The sparsity and Hessian continuity priors suppress this residual background while preserving weak vessel signals; consistently, the combined formulation retained the largest deep-signal fraction of all tested methods, whereas the sparsity prior alone tended to suppress weak deep vessels. This indicates that the continuity prior is important for maintaining deep sensitivity and that the two priors are complementary rather than redundant.

Compared with ultrasound localization microscopy [7,8], which provides micrometer-scale resolution but requires microbubble injection, acquisitions of several minutes, and dedicated post-processing, the present approach yields a more modest, several-fold narrowing of the apparent vessel width and operates entirely without contrast agents and on much shorter acquisitions. The two methods are therefore complementary: sparse deconvolution targets fast, contrast-free imaging where ULM cannot readily be deployed (e.g., bedside monitoring, repeated longitudinal studies, or contraindications to contrast agents), while ULM remains the method of choice when sub-PSF resolution is essential and microbubble injection is acceptable.

Beyond structural imaging, the same framework improved task-evoked functional ultrasound. Conventional functional ultrasound infers neuronal activity from stimulus-locked changes in the power-Doppler signal, but its spatial resolution is bounded by the same point-spread function that limits the underlying microvascular image, so that activation is reported as a diffuse cortical region rather than resolved to individual vessels. Applying sparse deconvolution to the power-Doppler time series preserved the stimulus-locked haemodynamic response—the conventional and deconvolved region-of-interest time courses were closely correlated—while confining the correlation-based activation map to discrete penetrating vessels, thereby increasing the spatial specificity of the functional map without any contrast agent or change of acquisition. Two points qualify this result. First, because the deconvolution is nonlinear, the reconstructed ΔCBV/CBV₀ amplitude is not a calibrated haemodynamic measure; quantitative response amplitudes should be read from the conventional power Doppler, whereas sparse deconvolution contributes the vessel-level localization. The activation maps themselves, being derived from a correlation with the stimulus, are invariant to monotonic intensity transformations and therefore remain a fair basis for comparing spatial specificity. Second, sparse deconvolution provides this vessel-scale localization without microbubbles, positioning it as a contrast-free complement to functional ULM, which resolves single-vessel haemodynamics but requires contrast injection and longer acquisitions.

Several limitations should be acknowledged. First, like all iterative deconvolution algorithms, sparse deconvolution depends on the quality of the input SNR and on the choice of the fidelity and sparsity weights. Second, the FWHM and the related sparsity metrics reflect the apparent width of the reconstructed vessels and the concentration of intensity, not a calibrated point-spread resolution; because the sparsity prior actively concentrates intensity, the smallest in vivo FWHM values fall below the physical diffraction limit and must be read as comparative sharpness measures rather than as evidence of true sub-wavelength resolution. Third, the sparse-deconvolution activation maps are visibly sparser than their conventional counterparts, and it remains an open question whether this sparsification removes only point-spread-related blur or also discards genuine but weakly activated vascular compartments — for instance capillary-level or small venular contributions whose amplitude falls below the effective sparsity threshold. Systematically quantifying how the activated area depends on the sparsity weight, and validating it against a co-registered functional ULM or optical reference acquired on the same plane, is therefore an important direction for future work.

## 5. Conclusion

We have presented a sparse deconvolution framework for contrast-free microvascular imaging in ultrafast power Doppler ultrasound. By combining a Split-Bregman optimization with sparsity and Hessian continuity priors and an accelerated Richardson–Lucy deconvolution with a calibrated system PSF, the pipeline recovers contrast and sharpness from SVD-filtered Doppler images. On a numerical phantom with known ground truth, all sparse-deconvolution variants reduced the reconstructed vessel FWHM relative to conventional SVD, with the combined sparsity-with- continuity formulation giving the finest apparent vessel width. In vivo experiments on the rat brain with a 15-MHz linear array showed increased sparsity and dynamic range, a several-fold narrowing of the apparent vessel width, and extended spatial-frequency content, while the combined formulation additionally preserved deep-vessel signal and a velocity-band decomposition disentangled slow and fast flow into a velocity-resolved map. Applied to task-evoked functional ultrasound, the framework preserved the stimulus-evoked cerebral-blood-volume response while localizing the activation from a diffuse cortical region to discrete penetrating vessels, extending contrast-free sparse deconvolution from structural to functional imaging.

## Acknowledgements

This work was supported by the Inserm research accelerator (Inserm ART) in Biomedical Ultrasound and by the French National Research Agency (ANR) under the ANR-21-CE19-0050 program (Project SonoGT). This research was also funded by the Region Ile de France— Convention DIM ELICIT Innovative Technologies for Life Science, and partially funded by the AXA Research Fund (Project NeuroElastoFlow).

## Competing Interests

The authors declare that the research was conducted in the absence of any commercial or financial relationships that could be construed as a potential conflict of interest.

## Supplementary Material

The following analysis supports, but is not required by, the main conclusions of the paper. It provides a data-driven alternative to the supervised, regressor-based activation mapping reported in Section 3.5.

### S1. Regressor-free extraction of the functional response by singular value decomposition

The correlation-based activation maps of Section 3.5 are supervised: they require the stimulus timing to be known so that a regressor can be correlated against every pixel time course. We reasoned that, because the whisker-stimulation paradigm imposes a strong periodic modulation on the power-Doppler time series, the stimulus-related component should emerge automatically as one of the dominant modes of a singular value decomposition (SVD) of the functional recording, without any prior knowledge of the stimulus timing. Decomposing the block time series by SVD factorizes it into paired spatial and temporal singular vectors ranked by the variance they explain; a mode whose temporal singular vector oscillates at the stimulation frequency then identifies the task-related spatial pattern in a fully data-driven manner.

The block design used here — 40 s of rest followed by 30 s of stimulation, repeated over five cycles — has a fundamental period of 70 s and hence a characteristic frequency of approximately 0.015 Hz. Taking the Fourier transform of each temporal singular vector and inspecting the resulting spectra (Fig. S1(a)) showed that the stimulus frequency was concentrated in a single low- order mode for each dataset: the fifth temporal singular vector for the conventional power-Doppler dataset and the fourth for the sparse-deconvolution dataset. The remaining low-order modes captured slow physiological drift and global haemodynamic fluctuations at frequencies distinct from the stimulus, and were readily separated from the task-related mode on the basis of their temporal spectra.

The spatial singular vectors associated with these task-related modes reproduced the activation pattern obtained by supervised correlation mapping. Among the first eight spatial singular vectors of each dataset (Fig. S1(b)), the identified mode exhibited a spatial structure co-localized with the barrel-cortex activation, while the higher-variance modes captured the bulk vascular anatomy and clutter residuals. Converting the corresponding temporal singular vector back to the time domain showed that it followed the block-design envelope and was statistically indistinguishable from the supervised stimulus-locked response (non-significant difference; Fig. S1(c)), confirming that the SVD recovers the same task-evoked signal that the correlation analysis extracts with explicit knowledge of the paradigm.

Notably, the task-related mode appeared at a lower rank in the sparse-deconvolution dataset than in the conventional one (fourth versus fifth singular vector). Because SVD orders modes by explained variance, this shift indicates that sparse deconvolution concentrates a larger share of the stimulus-related variance into a single, more dominant component, consistent with its suppression of the diffuse background and its confinement of the response to a compact set of vessels, which together reduce the mixing of the task signal with clutter. These results establish that SVD provides a label-free, regressor-free route to the functional activation map, and that sparse deconvolution renders the task-related component more separable, making automatic detection of the activated vasculature more robust.

**Figure S1.**
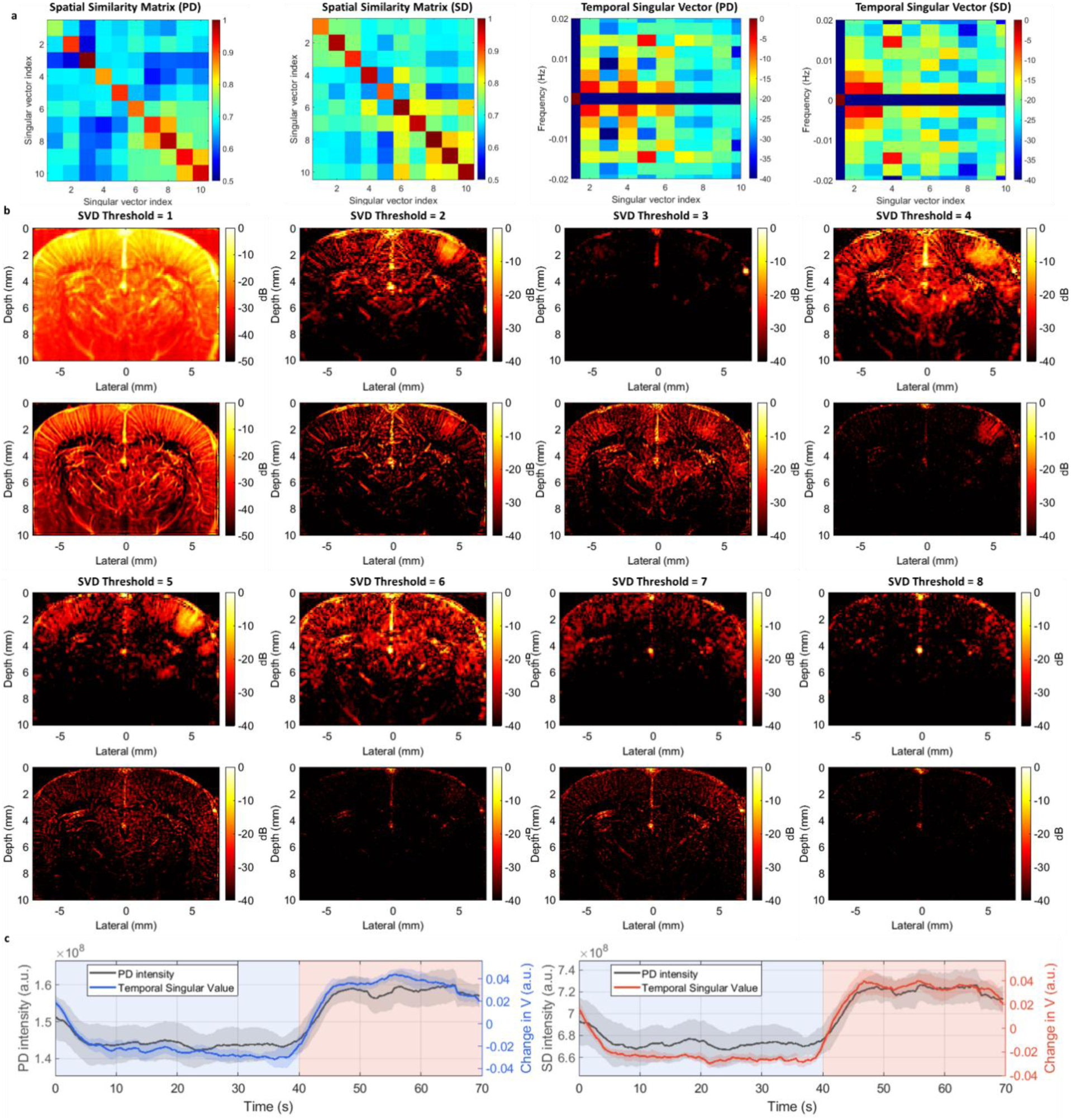
Data-driven extraction of the functional response by singular value decomposition (SVD) of the power- Doppler time series. (a) Spatial similarity matrix identifying the stimulus-related mode: the matrix relates the spatial singular vectors to the correlation-based activation pattern, while the Fourier transform of the temporal singular vectors isolates the mode whose temporal spectrum peaks at the stimulation frequency (≈0.015 Hz) — the fifth mode for the PD dataset and the fourth for the SD dataset. (b) First eight spatial singular vectors of the PD and SD datasets; the stimulus-related mode (identified in (a)) is spatially co-localized with the barrel-cortex activation, whereas the higher-variance modes capture the bulk vascular anatomy and clutter residuals. (c) Temporal singular vector of the identified mode converted back to the time domain, following the block-design envelope and showing a non- significant difference from the stimulus-locked activation obtained by correlation mapping.

## Notes

### Competing Interest Statement

The authors have declared no competing interest.

## References

1. Demené, C., et al., Spatiotemporal Clutter Filtering of Ultrafast Ultrasound Data Highly Increases Doppler and fUltrasound Sensitivity. IEEE Trans Med Imaging, 2015. 34(11): p. 2271–85.

2. Baranger, J., et al., Adaptive Spatiotemporal SVD Clutter Filtering for Ultrafast Doppler Imaging Using Similarity of Spatial Singular Vectors. IEEE Trans Med Imaging, 2018. 37(7): p. 1574–1586.

3. Mace, E., et al., Functional ultrasound imaging of the brain: theory and basic principles. IEEE Trans Ultrason Ferroelectr Freq Control, 2013. 60(3): p. 492–506.

4. Demené, C., et al., 4D microvascular imaging based on ultrafast Doppler tomography. Neuroimage, 2016. 127: p. 472–483.

5. Osmanski, B.F., et al., Functional ultrasound imaging of intrinsic connectivity in the living rat brain with high spatiotemporal resolution. Nat Commun, 2014. 5: p. 5023.

6. Macé, É., et al., Whole-Brain Functional Ultrasound Imaging Reveals Brain Modules for Visuomotor Integration. Neuron, 2018. 100(5): p. 1241–1251.e7.

7. Errico, C., et al., Ultrafast ultrasound localization microscopy for deep super-resolution vascular imaging. Nature, 2015. 527(7579): p. 499–502.

8. Couture, O., et al., Ultrasound Localization Microscopy and Super-Resolution: A State of the Art. IEEE Trans Ultrason Ferroelectr Freq Control, 2018. 65(8): p. 1304–1320.

9. Zhao, W., et al., Sparse deconvolution improves the resolution of live-cell super-resolution fluorescence microscopy. Nature Biotechnology, 2022. 40(4): p. 606–617.

10. Chen, Z., A. Basarab, and D. Kouamé, Compressive Deconvolution in Medical Ultrasound Imaging. IEEE Trans Med Imaging, 2016. 35(3): p. 728–37.

11. Michailovich, O. and A. Tannenbaum, Blind deconvolution of medical ultrasound images: a parametric inverse filtering approach. IEEE Trans Image Process, 2007. 16(12): p. 3005–19.

12. Jensen, J.A., Deconvolution of ultrasound images. Ultrasonic Imaging, 1992. 14(1): p. 1–15.

13. Bar-Zion, A., et al., *SUSHI: Sparsity-Based Ultrasound Super-Resolution Hemodynamic Imaging.* IEEE Transactions on Ultrasonics, Ferroelectrics, and Frequency Control, 2018. 65(12): p. 2365–2380.

14. Kim, J., et al., Compressed Sensing-Based Super-Resolution Ultrasound Imaging for Faster Acquisition and High Quality Images. IEEE Transactions on Biomedical Engineering, 2021. 68(11): p. 3317–3326.

15. You, Q., et al., Contrast-Free Super-Resolution Power Doppler (CS-PD) Based on Deep Neural Networks. IEEE Trans Ultrason Ferroelectr Freq Control, 2023. 70(10): p. 1355–1368.

16. Sloun, R.J.G.v., et al., Super-Resolution Ultrasound Localization Microscopy Through Deep Learning. IEEE Transactions on Medical Imaging, 2021. 40(3): p. 829–839.

17. Arendt Jensen, J., et al., Super-Resolution Ultrasound Imaging Using the Erythrocytes- Part I: Density Images. IEEE Trans Ultrason Ferroelectr Freq Control, 2024. 71(8): p. 925–944.

18. Kou, Z., et al., High-Resolution Power Doppler Using Null Subtraction Imaging. IEEE Trans Med Imaging, 2024. 43(9): p. 3060–3071.

19. Zhao, W., et al., High-throughput 3D super-resolution ultrasound imaging. bioRxiv, 2025: p. 2025.08.20.671215.doi: 10.1101/2025.08.20.671215.

20. Montaldo, G., et al., Coherent plane-wave compounding for very high frame rate ultrasonography and transient elastography. IEEE Trans Ultrason Ferroelectr Freq Control, 2009. 56(3): p. 489–506.

21. Richardson, W.H., Bayesian-Based Iterative Method of Image Restoration*. Journal of the Optical Society of America, 1972. 62(1): p. 55–59.

22. Biggs, D.S.C. and M. Andrews, Acceleration of iterative image restoration algorithms. Applied Optics, 1997. 36(8): p. 1766–1775.

23. Renaudin, N., et al., Functional ultrasound localization microscopy reveals brain-wide neurovascular activity on a microscopic scale. Nature Methods, 2022. 19(8): p. 1004–1012.

